# Structure of maize BZR1-type β-amylase BAM8 provides new insights into its noncatalytic adaptation

**DOI:** 10.1101/2022.05.09.491193

**Authors:** Fuai Sun, Malathy Palayam, Nitzan Shabek

## Abstract

Plant β-Amylase (BAM) proteins play an essential role in growth, development, stress response, and hormone regulation. Despite their typical (β/α)^8^ barrel structure as active catalysts in starch breakdown, catalytically inactive BAMs are implicated in diverse yet elusive functions in plants. The noncatalytic BAM7/8 contain N-terminal BZR1 domains and were shown to be involved in the regulation of brassinosteroid signaling and possibly serve as sensors of yet an uncharacterized metabolic signal. While the structures of several catalytically active BAMs have been reported, structural characterization of the catalytically inactive BZR1-type BAMs remain unknown. Here, we determine the crystal structure of *Zea mays* BZR1-type BAM8 and provide comprehensive insights into its noncatalytic adaptation. Using structural-guided comparison combined with biochemical analysis and molecular dynamics simulations, we revealed conformational changes in multiple distinct highly conserved regions resulting in rearrangement of the binding pocket. Altogether, this study adds a new layer of understanding to starch breakdown mechanism and elucidates the acquired adjustments of noncatalytic BZR1-type BAMs as putative regulatory domains and/or metabolic sensors in plants.

## Introduction

Plant β-amylases (BAMs) are members of the glycosyl hydrolase (GH)-14 family, which hydrolyze starch into maltose by cleaving α-1,4 glycosidic linkage from the non-reducing end of amylose (Totsuka and Fukazawa, 1996; Fulton et al., 2008). Plant genomes encode multiple β-amylases, however, not all are found to be catalytic enzymes (Thalmann et al., 2019). Most BAMs are well characterized in *Arabidopsis* (At), which is reported to have nine members of BAMs (BAM1-9) (Monroe et al., 2017) with conserved catalytic domains and variable N-terminal regions involved in localization. Among nine distinctive members of AtBAMs, AtBAM1/2/3/5 encode catalytically active enzymes (Monroe et al., 2017; Monroe et al., 2018) whereas the other three AtBAM4/7/8 are catalytically inactive. The sequence conservation analysis of these nine BAMs indicates that none of them share more than 60% homology, suggesting that they have not evolved recently through gene duplication events (Fulton et al., 2008). Interestingly, AtBAM1/2/3/4/6 are mainly localized in plastids where the starch is largely reserved. However, only AtBAM1/2/3 were reported as catalytically active enzymes. The distinct localization of BAMs suggests more diverse functions in addition to previous characterized catalytic activity (Lao et al., 1999; Fulton et al., 2008; Zybailov et al., 2008). AtBAM7 and AtBAM8 are members of BZR1 (brassinazole-resistant1)-type BAM family, which localized in the nucleus and have an additional BZR1 N-terminal DNA binding domain (Reinhold et al., 2011; Soyk et al., 2014). It has been proposed that AtBAM7/8 lack the starch hydrolysis activity and serve as transcriptional regulators in brassinosteroid (BR) signaling pathway (Reinhold et al., 2011). In corn, ZmBES1/BZR1-5 (*Zea mays*, BZR1-type BAM8) was reported to regulate plant development and interact with other proteins via BAM and/or BZR1 domains (Sun et al., 2020; Sun et al., 2021). Notably, several crystal structures of catalytically active plant BAMs have been determined from soybean (Mikami et al., 1993; Mikami et al., 1994), sweet potato (Cheong et al., 1995), barley (Mikami et al., 1999). Despite the advances in understanding their physiological roles in plants, the structure and function of noncatalytic BZR1-type BAMs remain unknown.

In this study, we determined the crystal structure of the BAM8 domain of ZmBES1/BZR1-5 (named ZmBZR1^BAM8^) and provided the detailed structural insights into its noncatalytic adaptation. Using structural-guided comparison combined with biochemical analysis and molecular dynamics simulations, we revealed conformational changes in multiple distinct highly conserved regions including the inner loop, the flexible loop, and a newly identified disulfide bridge. These structural modifications in ZmBZR1^BAM8^ result in rearrangement of the binding pocket, thus disrupting the perception and hydrolysis of linear saccharides. Altogether, this study adds a new layer of understanding of starch breakdown mechanism and elucidates the structural adjustments of noncatalytic BZR1-type BAMs as putative regulatory domains and/or metabolic sensors in plants.

## Results

### Overall Structure of ZmBZR1^BAM8^

To examine the BZR1-type BAM8 structure, we expressed and purified the β-amylase domain of ZmBES1/BZR1-5 (residues 203-651, termed ZmBZR1^BAM8^) and obtained protein crystals of the ZmBZR1^BAM8^ (**Fig. S1A**). The crystal structure of ZmBZR1^BAM8^ has been determined at 1.8Å resolution as a monomer in the asymmetric unit (**Table. 1**). ZmBZR1^BAM8^ consists of two structurally defined subdomains A and B. The central beta (β)_8_ strands in subdomain A are connected to form a barrel-shaped architecture. Each of the beta-strands in a barrel is surrounded by alpha (α)_8_ helices which together form the large (β/α)_8_ core region. The loops named L3 (residues 293-355), L4 (residues 384-460), and L5 (residues 497-525) are part of subdomain B, forming a smaller lobe of the (β/α)_8_ core region (**Fig. 1A** and **Fig. S1B**). Five short 3_10_ helices are found in which the first and last are present at the N- and C-terminal regions respectively whereas the other three 3_10_ helices are distributed in the smaller lobe of subdomain B. The putative catalytic site is located on a pocket-like cavity formed by distinct loops from subdomain A involving L6 (residues 544-561, contains the inner loop at positions 541-545), L7 (residue 586-591), and short loops from subdomain B involving L3 (contains the flexible loop at positions 298-304), L4, and L5 (**Fig. 1B-C**). This typical spatial arrangement of ligand-binding cavity is known to play role in the specific recognition of non-reducing end of linear sugar molecules (Davies and Henrissat, 1995). Our structural analysis shows that ZmBZR1^BAM8^ retained two conserved glutamic acid residues E388 and E584 that may serve as a nucleophile acceptor and a proton donor, respectively, as part of the catalytic process (**Fig. 1C** and **Fig. S1B**). Strikingly, the structure of ZmBZR1^BAM8^ reveals a unique disulfide bridge (S-S bond) between cysteine residues C589 and C630. This cysteine disulfide bridge is highly conserved among plant BAM8 but is missing in other BAM family members (**Fig. 1C, Fig. S1B, Fig. S2A-B**, and **Fig. S3**).

**Figure 1.**
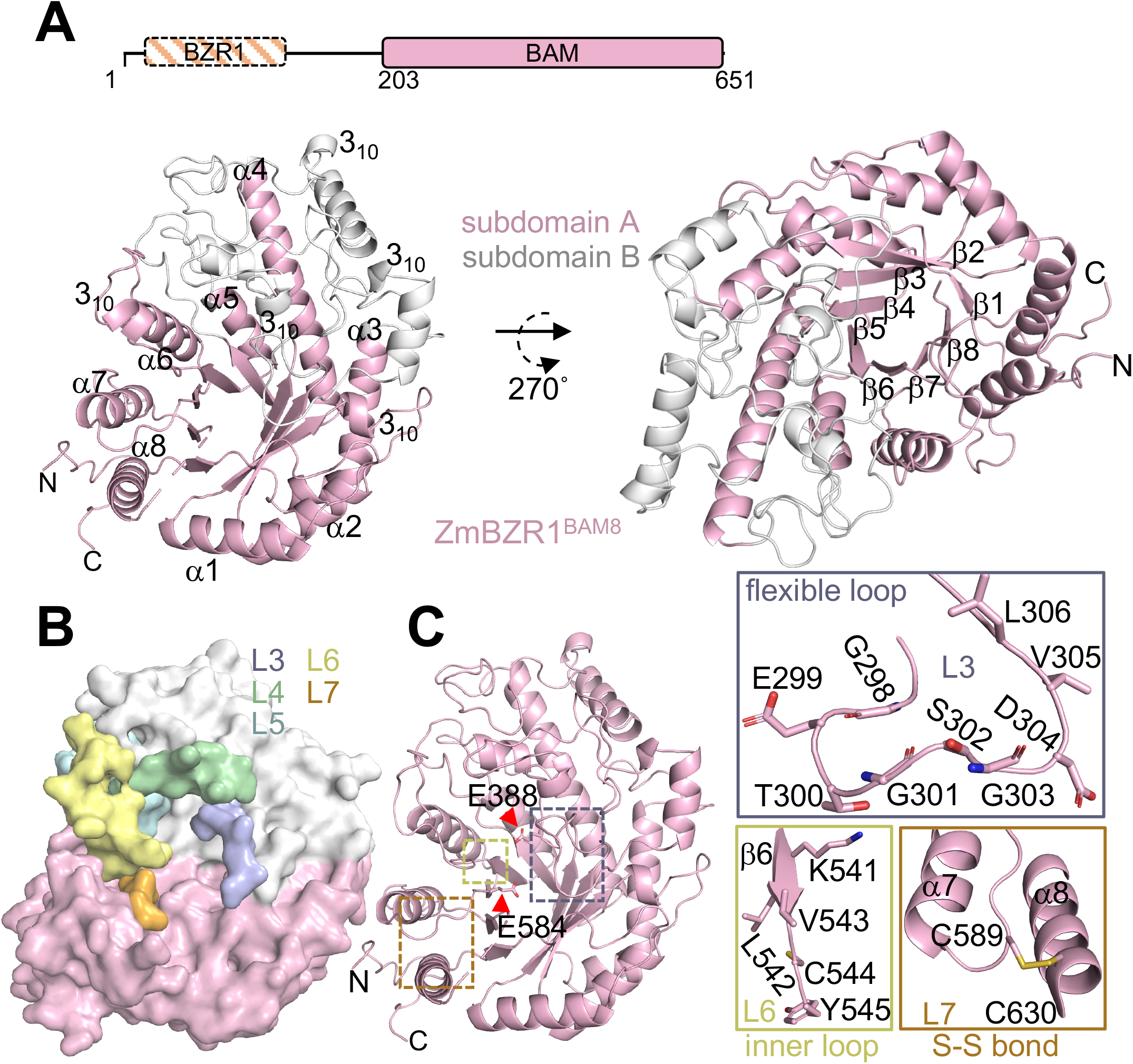
Molecular architecture of ZmBZR1^BAM8^. **A**. *Top*, Schematic representation of ZmBES1/BZR1-5 protein. *Bottom*, overall structure of ZmBZR1^BAM8^ is shown in side (*left*) and top views (*right*). Secondary structure of the larger core (β/α)_8_ subdomain A (pink) and smaller lobe, subdomain B (gray) represented as cartoon. **B**. Overall structure of ZmBZR1^BAM8^ shown in top view surface representation. Highlighted loops L3 (purple), L4 (green), L5 (cyan) L6 (yellow), and L7 (orange) are involved in forming the catalytic pocket. **C**. Close-up views of the flexible loop (*upper panel*), inner loop (*lower left*), and cysteine disulfide bridge (S-S bond, *lower right*) are shown in sticks and cartoons. The catalytic residues E388 and E584 are represented in sticks and indicated by red arrows.

### ZmBZR1^BAM8^ exhibits noncatalytic function

To further study the structure and function of the ZmBZR1-type BAM8, we attempted to co-crystallize and carry out serial crystal soaking experiments with increasing concentrations of starch or maltose as shown previously for GmBAM5-maltotetraose/maltose bound crystal structures (Kang et al., 2004). However, none of the newly resolved crystal structures contained any density of maltose despite the high-resolution electron densities in or around the catalytic pocket. We next sought to examine the catalytic function of ZmBZR1^BAM8^ by employing an optimized starch hydrolysis-based assay under various conditions (**Fig. 2A**). We first tested the effect of different pH conditions on starch hydrolysis activity. This effect has been previously reported for GmBAM5 where the pH optimum is around 5.4 due to the pKa of the amino acids within the catalytic pocket (Hirata et al., 2004). To that end, the enzymatic activity of ZmBZR1^BAM8^ was monitored by measuring the fluorescein release upon cleavage of saccharides in the presence of increasing pH conditions (from 3.66 to 8.11) (**Fig. 2A-B**). However, our data shows no catalytic activity for ZmBZR1^BAM8^ despite the changes in pH which aligns with our co-crystallization trials at pH 5.0.

**Figure 2.**
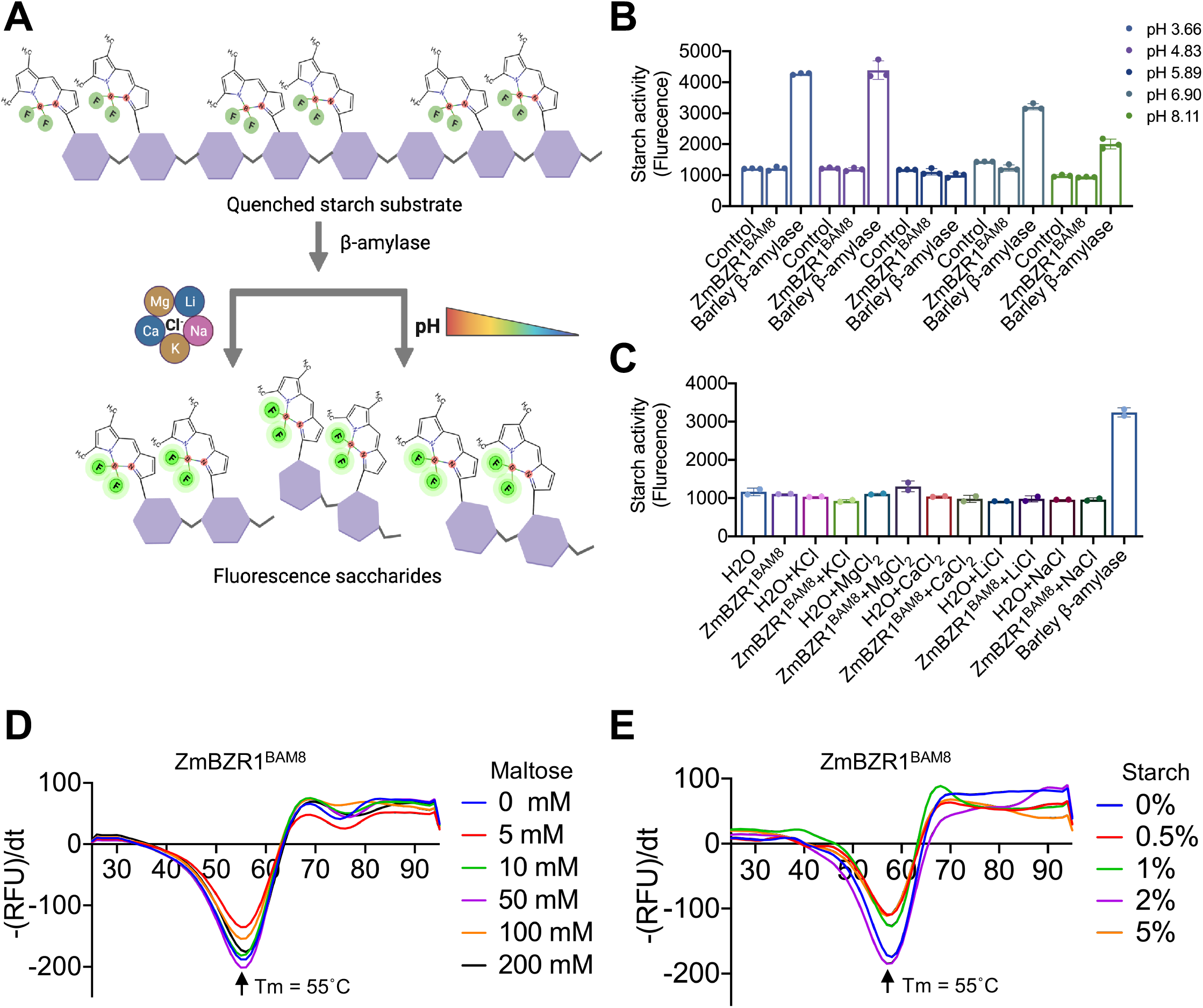
Inspecting ZmBZR1^BAM8^ enzymatic activity. **A**. Schematic representation of the amylase activity assay using fluorogenic starch as substrate. Generated by Biorender.com, 2022. **B-C**. Amylase activity assay of ZmBZR1^BAM8^ at the indicated pH conditions **(B)** or specified mono/divalent ions **(C)**. Barley β-amylase serves as a positive control. Data are means ± SE (error bar) with replicates (shown as circles). **D-E**. Differential Scanning Fluorimetry (DSF) analyses of ZmBZR1^BAM8^ in the presence of increasing concentrations of maltose **(D)** or starch **(E)**. (RFU, Related Fluorescence Units)/dt indicates the changes in fluorescence level per changes in temperature (25-95°C). Tm denoted melting temperature peak of ZmBZR1^BAM8^.

It has been previously shown that AtBAM2 activity is likely elevated in the presence of monovalent ions by increasing ionic strength of the assay (Monroe et al., 2017). To address this, we tested the activity of ZmBZR1^BAM8^ with monovalent ions such as KCl, NaCl, LiCl and divalent-containing salts (MgCl_2_ and CaCl_2_) (**Fig. 2C**), yet ZmBZR1^BAM8^ remains inactive. To overrule the possibility that ZmBZR1^BAM8^ can perceive sugar molecules without necessarily hydrolyzing it, we performed a thermal shift assay (Different Scanning Fluorimetry, DSF) with varying concentrations of maltose or starch. As expected, ZmBZR1^BAM8^ thermal stability remains intact even in very high concentrations of starch or maltose (**Fig. 2D-E**). Altogether, our data suggest ZmBZR1^BAM8^ has no catalytic activity compared to other BAMs which is consistent with the previous study of AtBAM8 (Soyk et al., 2014).

### Phylogenetic analysis of ZmBZR1^BAM8^

To investigate the evolution of catalytic and noncatalytic BAMs in plants, we retrieved the sequences of BAMs from different plants using the catalytic AtBAM1/2/3/5 and putative noncatalytic AtBAM7/8 as the query sequences. The phylogenetic analyses identified three distinct clades: the catalytic AtBAM1/3/5, the noncatalytic BAM7, and BAM8 as a separate group (**Fig. 3A**). Notably, the AtBAM2 falls into the same clades as noncatalytic BAM7 even though it is catalytically active and depends on the presence of a potassium ion (Monroe et al., 2017). Since the flexible and inner loops (**Fig. 1C** and **Fig. S1B**) were previously shown to play a crucial role in β-amylase enzymatic activity, we analyzed the conservation of these loops in all flowering plants. We found that both loops are highly conserved in the catalytic BAM1/2/3 and BAM5, whereas the flexible loop is conserved only in the noncatalytic BAM7 (**Fig. 3B**). The flexible and inner loops were not found to be highly conserved in all BAM8 orthologs, which is consistent with the previously reported noncatalytic function of BAM7 and BAM8 in Arabidopsis (Soyk et al., 2014). Interestingly, the highly conserved catalytic residues E388 and E584 (E186 and E380 in GmBAM5, respectively) were also found in the noncatalytic BAMs, corroborating the notion that other residues and structural rearrangements may play role in effective catalysis. Importantly, while the glutamine residue (Q94 in BAM1/3/5) preceding the flexible loop is conserved in the catalytic BAM1/3 and BAM5, it is altered to glutamic acid in the noncatalytic BAM7/8. Interestingly, at this position, ZmBZR1^BAM8^ which falls into the BAM8 subclades contains glycine (G296, **Fig. 3A-B**). Our analysis proposes that G296 preceding the flexible loop has been co-diverged with E299 within the flexible loop. We further identified a salt bridge formation between E299-N333 in ZmBZR1^BAM8^ crystal structure that stabilizes the flexible loop (**Fig. 4A, Fig S1B**). Our sequence conservation analysis proposes a structural adaptation in distinct regions in BAMs that may explain their catalytic and noncatalytic functions.

**Figure 3.**
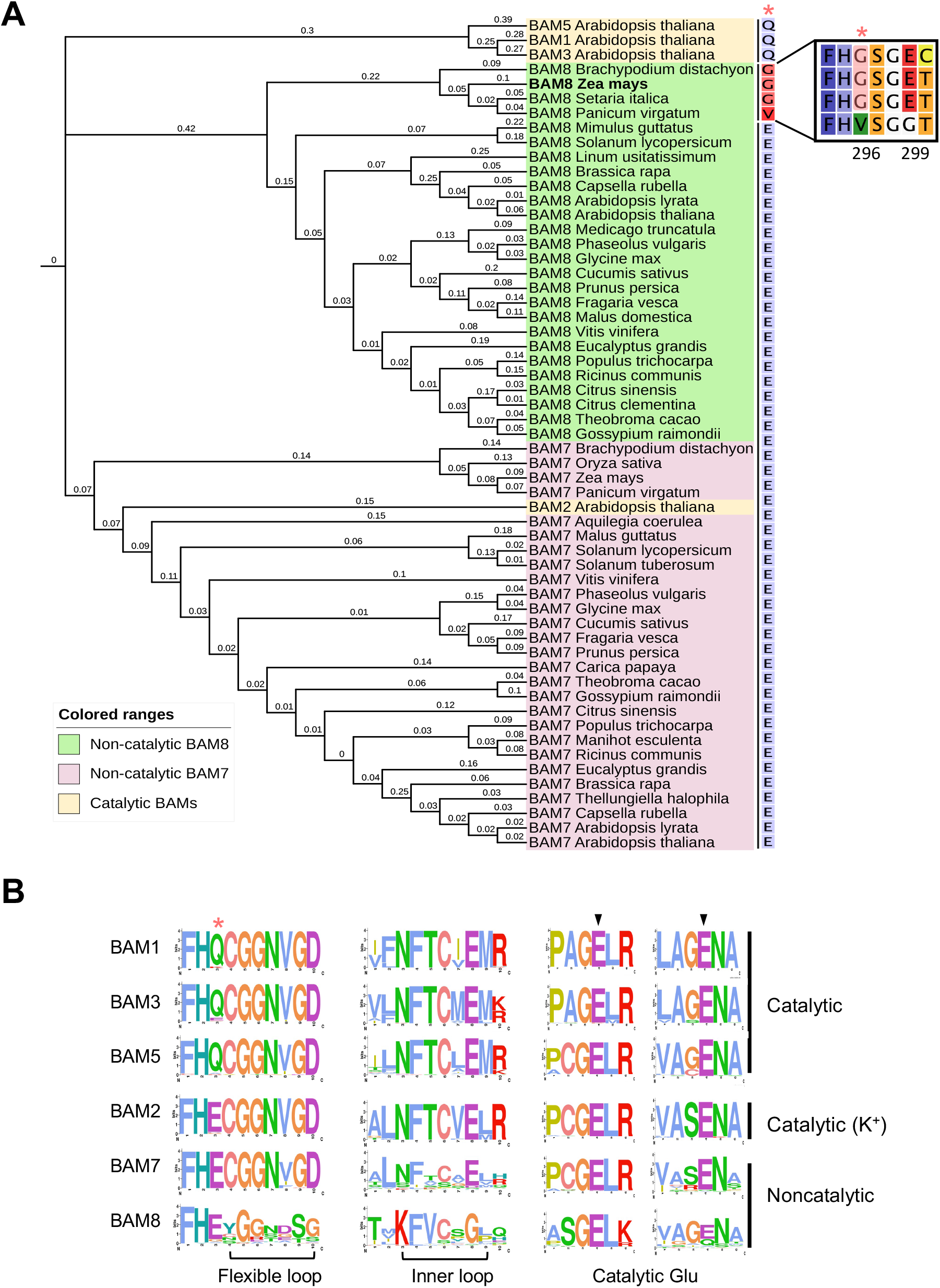
The phylogenetic and sequence conservation analyses of ZmBZR1^BAM8^. **A**. Phylogenetic analysis of catalytic BAM1/2/3/5, and noncatalytic BAM7/8 marked with yellow, pink, and green, respectively. The branch length is labeled and represents the measurement of substitutions per site. The framed square (*top right*) represents an extended sequence (294-300) of the indicated subclades. The single amino acid at the right side of the phylogenetic tree represents a residue preceding the flexible loop (red asterisk). **B**. The sequence logos of flexible loop, inner loop, and catalytic Glu are shown. The bit score indicates the information content of each position. The relative size of the amino acids indicates their conservation in the sequence. All sequences and species accession numbers are listed under *Supplementary Datasets 1 and 2*.

**Figure 4.**
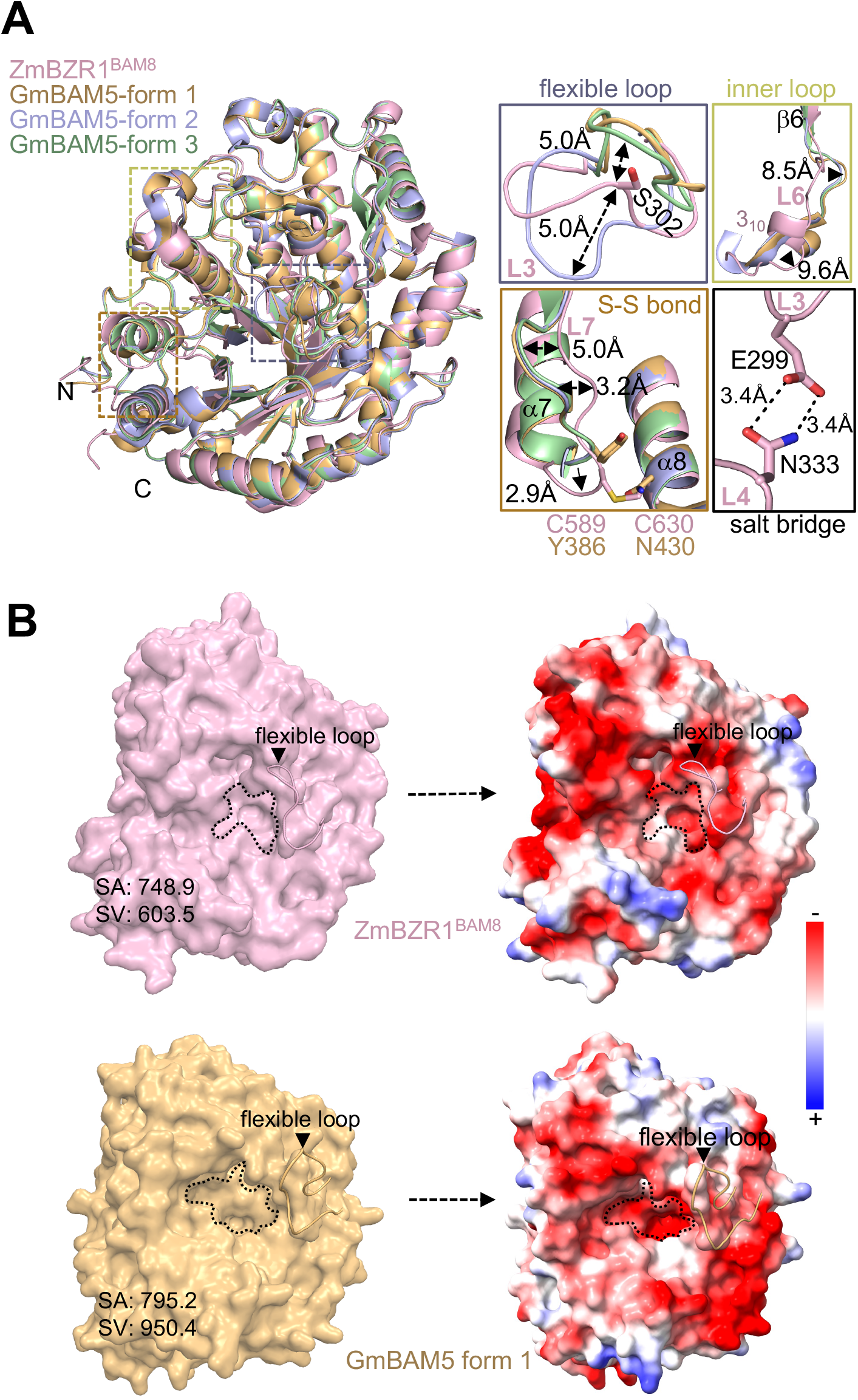
Comparative structural variations of ZmBZR1^BAM8^. **A**. Superposition of ZmBZR1^BAM8^ (light pink) with GmBAM5 shown in side view. The RMSD of 0.8Å, 0.7Å, and 0.7Å are calculated for superposition between ZmBZR1^BAM8^ and GmBAM5 (form 1: Apo, wheat color, PDB: 1WDP), GmBAM5 (form 2: post-hydrolysis, purple, PDB: 1Q6C) and GmBAM5 (form 3, product release: pale green, PDB: 1BTC) respectively. *Right upper panels*, close-up views of the flexible loop and inner loop of ZmBZR1^BAM8^ with GmBAM5. *Right lower panels*, close-up view of cysteine disulfide bridge (S-S bond) in stick representation *(left)* and residues involved in the salt bridge *(right)*. **B**. Comparison of electrostatic surface representation of ZmBZR1^BAM8^ (*top left*, light pink) and GmBAM5 (*bottom left*, wheat color). The active site pocket is highlighted in dashed lines. The electrostatic potential is calculated by PyMOL and APBS with the non-linear Poisson-Boltzmann equation contoured at ±5 *k*T/*e*. Negatively and positively charged surface areas are colored in red and blue, respectively.

### Structural comparative analysis of ZmBZR1^BAM8^ reveals alterations in spatial rearrangement of the binding pocket

Much of the knowledge about plant β-amylase function stems from the structural and biochemical studies of BAM5, yet a comprehensive understanding of other BAM orthologs, particularly the BZR1-type BAMs remain to be explored. Towards that end, we analyzed the molecular structure of ZmBZR1^BAM8^ by carrying out detailed structure-guided comparative characterization with the previously reported GmBAM5 structure. Despite the structural similarities, ZmBZR1^BAM8^ reveals significant structural and amino acid changes within the binding pocket and active sites (**Fig. 4A**). It has been previously shown that L3, containing the conserved flexible loop, can exist in two distinct conformations: an “open” state, that is solvent-exposed, and a “closed” state, where L3 shifts towards the substrate upon binding (Kang et al., 2005). This conformational shift has been implicated in saccharides perception as well as the release of the hydrolyzed products (Mikami et al., 1994; Kang et al., 2005). In GmBAM5, residues V99 and D101 within L3 directly interact with the sugar molecule via van der Waals and hydrogen bonds respectively. In ZmBZR1^BAM8^, the L3 region is found in a unique intermediate state between the “open” and “closed” conformation (**Fig. 4A** and **Fig. S2A**). A sharp kink is observed at S302 in ZmBZR1^BAM8^ leading to a substantial deviation (∼5.0Å) compared to the canonical open/closed states of other plant BAM5 reported structures (**Fig. 4A** and **Fig. S2A-B**). Moreover, the region of L3 in position 298-301 (GETG) appears to be pushed towards L4 in ZmBZR1^BAM8^ and stabilized by a salt bridge between E299 and N333 (E299-N333, **Fig. 4A**). Notably, the residue E299 in ZmBZR1^BAM8^ is completely replaced by G98 in BAM5, thus preventing stabilization by ionic interaction (**Fig. S2B**). The partial hydrophobic nature of L3 in BAM5, particularly the residues involved in ligand coordination, has been diverged into polar uncharged amino acids, such as T300 and S302 in ZmBZR1^BAM8^, resulting in a more polar flexible loop compared to the BAM5 (**Fig. 4B** and **Fig. S2B)**. These structural changes in L3 may affect the perception of sugar molecules by ZmBZR1^BAM8^ and explain its inactive state. This is consistent with the previously reported D101N mutant in GmBAM5 that shows no detectable catalytic activity (TOTSUKA et al., 1994).

The short inner loop segment in GmBAM5 plays the second major role in positioning the substrate by acting as a “trap-trigger” to appropriately orient the sugar molecule inside the catalytic cavity. The residues F-T-C (341-343) in the inner loop are highly conserved in BAM1/2/3/5 (**Fig. 3B** and **Fig. S2B)**. In GmBAM5, the key threonine residue, T342, found in two distinct orientations (apo and post-hydrolysis), interacts with catalytic E186 and sugar molecule substrates (Kang et al., 2004). In ZmBZR1^BAM8^, the polar T342 substituted to the hydrophobic V543 contributed to the increase in overall hydrophobicity of the inner loop (**Fig. 4B** and **Fig. S2B**). This substitution resulted in the loss of the hydrogen bond network with the catalytic glutamic acid, thus making it unable to stabilize its deprotonated state during the catalysis. It has been shown that T342V substitution in mutant GmBAM5 (GmBAM5_T342V_) has a 13-fold decrease in catalytic activity compared to the wildtype where the main chain oxygen atom of V342 forms a hydrogen bond with the glucose moiety (Kang et al., 2005). At the same position, the natural valine (V543) in ZmBZR1^BAM8^ is oriented farther away (6.3 Å) from the putative glucose moiety (O4, **Fig. S2C**) compared to GmBAM5_T342V_. Further analysis reveals that the inner loop in ZmBZR1^BAM8^ adopts a distinct conformation with a type II β-turn (positions 546-549) and 3_10_ helix (positions 550-552) that were missing in other BAMs crystal structures (**Fig. 4A, Fig. S1B** and **Fig. S2A**). This type II β-turn conformation results in ∼180° flip in the center of the inner loop. Furthermore, part of L7 in ZmBZR1^BAM8^ (residues 584-589, and 380-385 in GmBAM5) is unable to perceive the substrate as in GmBAM5. Instead, it is pushed away from L6 towards α8 on the opposite side. This movement is a direct result of a S-S bond between C589 of L7 and C630 of α8 in the ZmBZR1^BAM8^ structure (**Fig. 1C, Fig. 4A**, and **Fig. S2A-B**). Remarkably, these cysteines are uniquely conserved in BAM8 to accommodate the S-S bond. Additional analysis of other BAM orthologs shows sequence variations in the position of the S-S bond, for example, the cysteines are substituted with tyrosine (Y386) and asparagine (N430) in the BAM5 orthologs (**Fig. 4A, Fig. S2A-B**, and **Fig. S3**). The structures reported previously together with our current high-resolution structures, allow us to resolve altered orientations of distinct side chains that replace maltose contacts with water molecules in ZmBZR1^BAM8^ (**Fig. S4A-B**).

Taken together, our structural analysis explains the noncatalytic function of ZmBZR1^BAM8^ as well as its inability to effectively perceive the substrate despite the presence of the catalytic residues.

### Molecular dynamics simulations of ZmBZR1^BAM8^ reveal structural rigidity of the flexible and inner loops

To further study the plasticity of the flexible and inner loops in ZmBZR1^BAM8^, a comparative molecular dynamics (MD) simulation was performed using an AMBER99SB-ILDN force field for ZmBZR1^BAM8^ and GmBAM5 (Apo, form 1). The simulation trajectories were analyzed for the structural stability of the apo form of the enzyme by calculating the root mean square deviation (RMSD) and root mean square fluctuation (RMSF) over 100 nanoseconds simulation. The backbone RMSD values of both proteins increased during the initial phase of simulation after which a plateau was converged at 0.17 nm (**Fig. 5A**). The deviations around 50-60 ns in ZmBZR1^BAM8^ and 60-80 ns in GmBAM5 are likely due to the different conformation of the flexible (L3) and inner (L6) loop between the structures (**Fig. 5B**). Moreover, the flexible and inner loops in GmBAM5 are more dynamic than ZmBZR1^BAM8^ as can be seen by the RMSF for each structure (**Fig. 5B**). As expected, the S-S bond in ZmBZR1^BAM8^ exhibits higher dynamics due to the breaking and formation of the disulfide bridge during the simulation (**Fig. 5A-B**). Thus, our MD analysis aligns with the structural rigidity of ZmBZR1^BAM8^ as can be seen in our comparative analysis (**Fig. 5B-C**) and corroborates the requirements of the inner and flexible loops’ plasticity to effectively catalyze saccharides (**Supplementary Movie 1** and **Supplementary Movie 2**). This data further supports the inability to perceive and catalyze the by BZR1-type BAM8 in plants.

**Figure 5.**
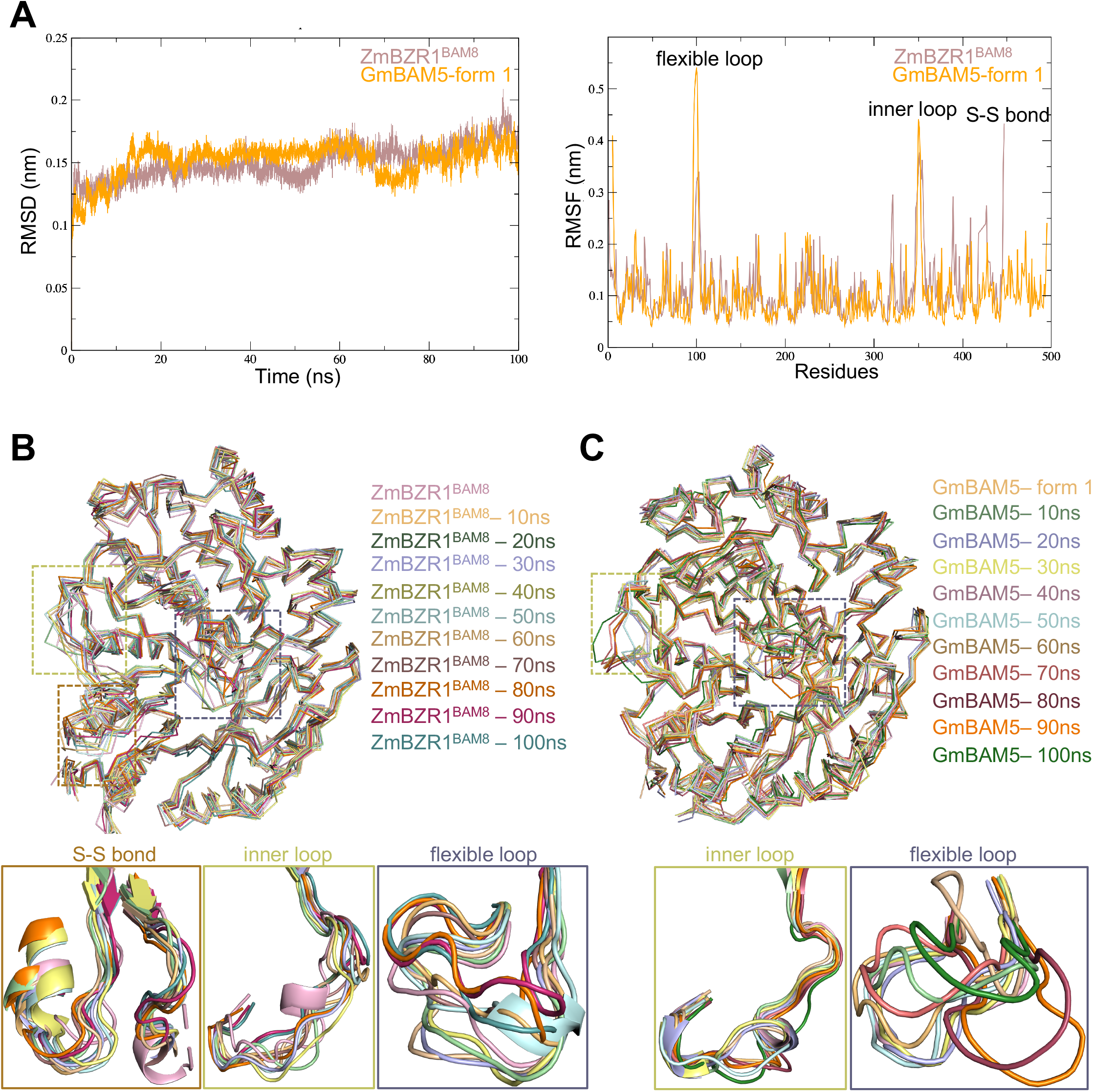
Comparative molecular dynamics simulation of ZmBZR1^BAM8^ and GmBAM5. **A**. (*left*) RMSD graphs plotted for the backbone atoms of ZmBZR1^BAM8^ (pink) and GmBAM5 (yellow, PDB: 1WDP). (*Right*) Root mean square fluctuation (RMSF) of ZmBZR1^BAM8^ and GmBAM5 versus residue number. **B**. Structural superposition of ZmBZR1^BAM8^ retrieved from different time points (*Top*). Close-up view of the cysteine disulfide bridge (S-S bond, *Bottom left panel*), the inner loop (*Bottom middle panel*), and the flexible loop (*Bottom right panel*), at different time points of the simulation are represented as cartoon. **C**. Structural superposition of GmBAM5 retrieved from different time points (*Top*). Close-up view of the inner loop (*Bottom left*) and flexible loop (*Bottom right*) at different time points of the simulation are represented as cartoon

## Discussion

β-amylase (BAM) plays an important role in starch degradation during the day-night cycle of plants (Ishihara et al., 2022). While BAMs have been extensively characterized in Arabidopsis (AtBAM1-9), there is limited knowledge of their functions in other plants. Interestingly, not all BAMs have been reported to function as catalytic amylases. For example, the noncatalytic AtBAM7/8 are members of the BZR1-type BAMs which have additional N-terminal BZR1 domains, as well as BAM domain with elusive role in transcriptional regulation (Soyk et al., 2014). In this study, we reveal a high-resolution crystal structure of *Zea mays* BZR1^BAM8^ with new structural elements that have not been previously reported. Our biochemical and structural comparative analyses coupled with the MD simulations reveal the structural adaptation and spatial rearrangement requirements for both the catalytic and noncatalytic BAMs. In both the catalytic and noncatalytic BAMs, the flexible loop (in L3), the inner loop (in L6), and L7 represent the key structural elements needed to determine successful substrate binding and hydrolysis. Our current model proposes that upon substrate binding, L3 undergoes a substantial conformational change to stabilize the substrate together with L6 and L7 (**Fig. 6Ai-ii**) and allow the glutamic acid catalytic residues (in L7) to cleave the nonreducing end of the sugar molecules (**Fig. 6Aiii**). However, in the noncatalytic BZR1-type BAMs, L3 is rigid and L6 undergoes a conformational change that disrupts the access of the substrate to the catalytic cavity (**Fig. 6Bi**). Moreover, L7 which contains the catalytic glutamic acid is pushed away from the active site due to the disulfide bridge (S-S bond) that is formed with α8 (**Fig. 6Bi-ii**). The function of Arabidopsis noncatalytic BAM in BZR1 has been suggested to serve as a critical element in the regulation of DNA recognition and transcription (Soyk et al., 2014). Therefore, it is possible that the conformational rearrangements of the highly conserved key loops and other structural elements may participate in protein-protein interaction or in protein-DNA regulation, and/or serve as the intermolecular interface with other domains. One intriguing function that we cannot exclude is the possibility that ZmBZR1^BAM8^ and its orthologs may represent new metabolic sensors of yet an unidentified sugar-like ligand(s). The primary function of these noncatalytic BAMs remains elusive and future studies in this direction may reveal new regulatory mechanisms of BZR1-type BAMs in plants.

**Figure 6.**
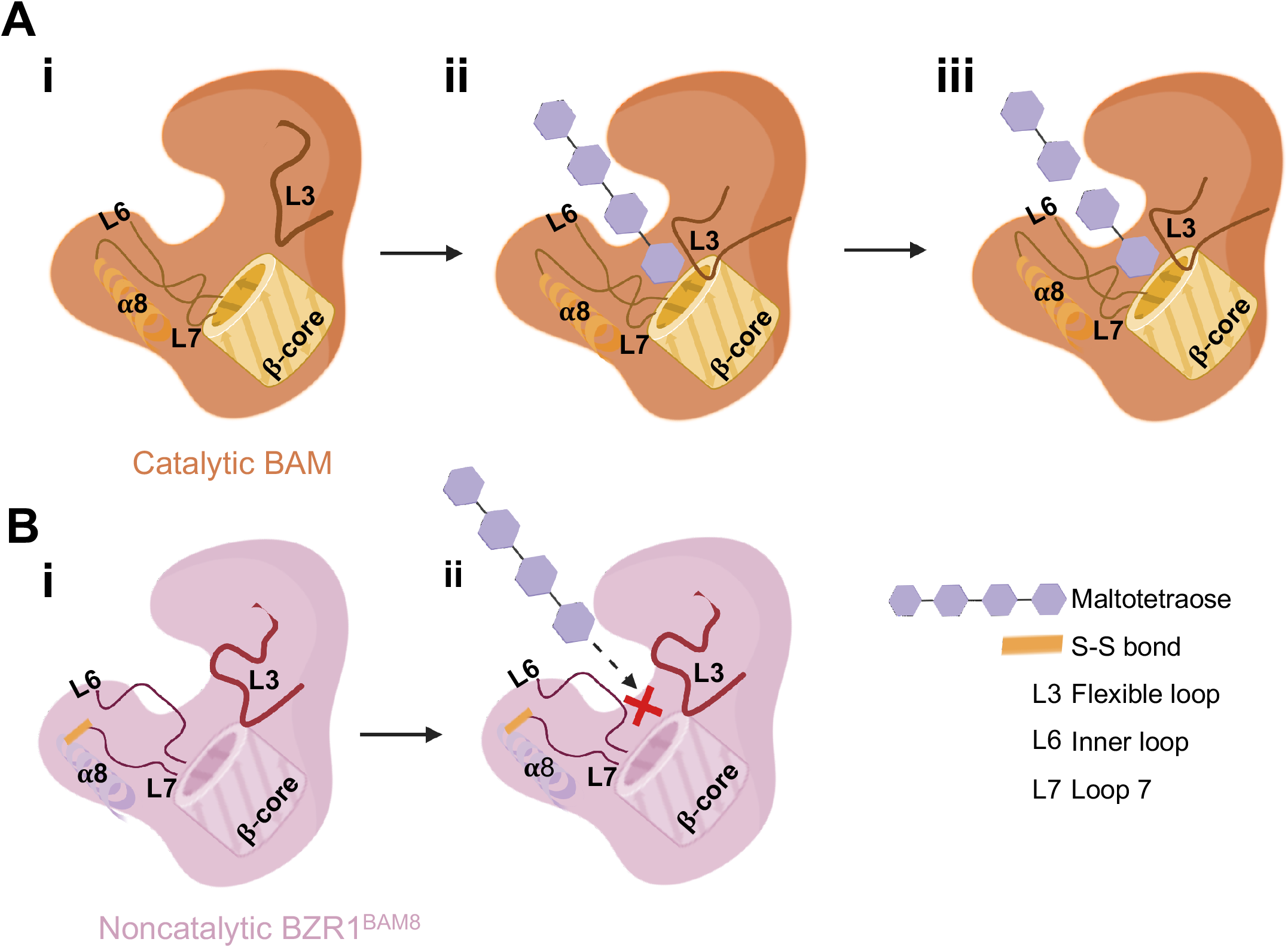
A proposed model of catalytic and noncatalytic BAMs. **A**. In the catalytic BAMs (**i-ii**) The flexible loop (L3) undergoes conformational changes when the substrate binds. The inner loop (L6) participates in positioning the substrate (polysaccharide chain, purple) together with loop (L7). **(iii)** The substrate is hydrolyzed by the catalytic residues in the loops L3, L6, and L7. **B. (i-ii)** In the noncatalytic BZR1^BAM8^, the flexible loops are positioned in an intermediate state (**i**) and the rearrangement of L6, and L7 together with the cysteine disulfide bridge (S-S bond) between L7 and α8 disrupts the both the active site and the binding pocket accessibility for the substrate **(ii)**. The graphical representation was generated using Biorender.com, 2022.

## Material and Methods

### Protein Expression and Purification

ZmBZR1^BAM8^ (ZmBES1/ZBR1-5, BAM in positions 203-651) was amplified with forward primer aaaacctctacttccaatcgATGTTATTCCCCGATGATTACACGA and reverse primer aaaacctctacttccaatcgATGTTATTCCCCGATGATTACACGA. The ZmBZR1^BAM8^ was expressed as 6x His-SUMO fusion protein from the pAL vector (Addgene). The verified plasmid was transformed into *Escherichia coli* strain BL21 (DE3), incubated at 37°C for 3-4 h until the OD_600_ reached 0.5, and then induced with 200 mM IPTG at 16°C overnight. Cells were harvested, resuspended, and lysed in extract buffer (20 mM Tris–HCl pH 8.0, 200 mM NaCl, 5mM 2-Mercaptoethanol, protease inhibitors). His-SUMO-ZmBZR1^BAM8^ was isolated from soluble cell lysate by Ni-NTA resin, and subsequently eluted with 200 mM imidazole. The eluates were subjected to anion exchange chromatography. The eluted protein was cleaved with TEV (tobacco etch virus) protease overnight at 4 °C. The cleaved His-SUMO was removed by passing tandem Ni-NTA and the purified ZmBZR1^BAM8^ protein was concentrated to ∼7 mg/ml for further crystallization and activity assays.

### Data Collection, Structure Determination, and Refinement

The ZmBZR1^BAM8^ protein crystallization experiments were performed by hanging-drop vapor-diffusion method at 25°C with a 2:1 ratio of protein and were equilibrated against a 2 ul reservoir containing 0.2 M MgCl_2_-6H_2_O, 0.1 M Sodium citrate tribasic dihydrate pH 5.0,8% w/v Polyethylene glycol 20,000. The crystals were harvested from a reservoir solution with 20% glycerol cryoprotectant and diffracted to 1.84 Å resolution; the data was collected and integrated, scaled by HKL 2000 (Otwinowski and Minor, 1997). The structure was solved by molecular replacement with GmBAM5 (PDB: 1WDP) as a searching model. The model was built and refined by PHENIX and COOT (Emsley and Cowtan, 2004; Afonine et al., 2018). All structures were generated by PyMOL (LLC., 2012). Atomic coordinates and structure factors for the reported crystal structure have been deposited in the Protein Data Bank with accession number 7UPV.

### Enzymatic Activity Assay and Different Scanning Fluorimetry (DSF)

ZmBZR1^BAM8^ activity was detected by EnzChek Ultra Amylase Assay Kit (Lot#E33651, Invitrogen, USA). The kit with 4,4-difluoro-3a,4a diaza-s-indacene dye (BODIPY FL)-labeled starch and relative fluorescence is proportional to amylase activity. Briefly, a total of 100 μL reaction mixture (done in triplicates) with different pH or salts were mixed with labeled starch using 96-well microplate, and barley β-amylase (A-7130, Sigma) served as a positive control. The fluorescence was detected by a microplate reader (Synergy, BioTek) set for excitation at 485 nm and emission at 530 nm.

DSF experiments were performed on the CFX96 Real-Time PCR system (Bio-Rad, Hercules, CA, USA). Sypro Orange was used as a reporter dye under excitation of 470 nm and emission of 575 nm (Life technology, Carlsbad, CA, USA). The PCR heat-denature with a gradient temperature from 25 - 95°C and final data were collected with CFX manager software. The reaction was carried out in a total of 30 μL comprises 20 μM ZmBZR1^BAM8^ protein, 0-200 mM maltose, and 0% - 5% starch in reaction buffer (20 mM HEPES buffer pH 7.3, 150 mM NaCl). All experiments were repeated at least three times and statistical analysis and graphical representations were performed by GraphPad Prism 8.0.

### Multiple Sequence Alignment and Phylogenetic Analyses

The β-amylase protein sequences were identified by BLASTp NCBI (https://www.ncbi.nlm.nih.gov/) using Arabidopsis catalytic BAM1/2/3/5 and noncatalytic BAM7/8 as the query sequence. The alignment of sequences was performed by MEGA11 using MUSCLE multiple alignment (Tamura et al., 2021). The phylogenetic tree was constructed by the Maximum Likelihood method (Jones et al., 1992) and colored with iDOL V5 (Letunic and Bork, 2021). The known conserved catalytic loop and residues were alignment by CLC Genomics Workbench v22 and the sequence logos were generated by WebLogo (Crooks et al., 2004) and TBtools (Chen et al., 2020). All sequences and species accession numbers are listed under *Supplementary Datasets 1 and 2*.

### Molecular Dynamics

Molecular dynamics (MD) simulations of ZmBZR1^BAM8^ and GmBAM5 (apo, form 1) were performed using GROMACS 2020.3 software package with AMBER99SB-ILDN force field (Davies and Henrissat, 1995) and the flexible SPC water model. The initial structure was immersed in a periodic water box of cubic shape (1.0 nm). Electrostatic energy was calculated using the particle mesh Ewald method (Wang et al., 2016), which permits the use of Ewald summation at a computational cost comparable to that of a simple truncation method of 10 Å or less. We retained the cutoff distance as 1.0 nm for the calculation of the coulomb and van der Waal’s interaction, respectively. After energy minimization using a steepest descent for 1000 steps, the system was subjected to equilibration at 300 k and normal pressure for 1000 ps. We subsequently applied LINCS (Hess, 2008) constraints for all bonds, keeping the whole protein molecule fixed and allowing only the water molecule to move to equilibrate with respect to the protein structure. The system was coupled to the external bath by the Berendsen pressure and temperature coupling (Eslami et al., 2010). The final MD calculations were performed for 100 ns under the same conditions except that the position restraints were removed. The results were analyzed using the software provided by the GROMACS package and graphs were plotted using Xmgrace V5.1.25.

## Author Contribution

N.S., F.S., and M.P designed and conceived the experiments. F.S. conducted the protein purification, crystallization, and biochemical assay. N.S. and M.P. determined and analyzed the structures. F.S., M.P., and N.S wrote the manuscript.

## Acknowledgments

N.S. is supported by NSF-CAREER (Award #2047396) and NSF-EAGER (Award #2028283). We thank the beamline staff at the Advanced Light Source (U.S. DOE Office of Science User Facility under Contract No. DE-AC02-05CH11231, is supported in part by the ALS-ENABLE program funded by the National Institutes of Health, National Institute of General Medical Sciences, grant P30 GM124169-01). F.S was partially supported by China Scholarship Council (CSC201906910056).

